# Three-Dimensional Label-Free Imaging and Quantification of Migrating Cells during Wound Healing

**DOI:** 10.1101/2020.07.24.219774

**Authors:** A. J. Lee, H. Hugonnet, W.S. Park, Y.K. Park

## Abstract

The wound healing assay provides essential information about collective cell migration and cell-to-cell interactions. It is a simple, effective, and widely used tool for observing the effect of numerous chemical treatments on wound healing speed. To perform and analyze a wound healing assay, various imaging techniques have been utilized. However, image acquisition and analysis are often limited in two-dimensional space or require the use of exogenous labeling agents. Here, we present a method for imaging large-scale wound healing assays in a label-free and volumetric manner using optical diffraction tomography (ODT). We performed quantitative high-resolution three-dimensional (3D) analysis of cell migration over a long period without difficulties such as photobleaching or phototoxicity. ODT enables the reconstruction of the refractive index (RI) tomogram of unlabeled cells, which provides both structural and biochemical information about the individual cell at subcellular resolution. Stitching multiple RI tomograms enables long-term (24 h) and large field-of-view imaging (> 800 × 400 μm^2^) with a lateral resolution of 110 nm. We demonstrated the thickness changes of leading cells and studied the effects of cytochalasin D. The 3D RI tomogram also revealed increased RI values in leading cells compared to lagging cells, suggesting the formation of a highly concentrated subcellular structure.

**STATEMENT OF SIGNIFICANCE:** The wound healing assay is a simple but effective tool for studying collective cell migration (CCM) that is widely used in biophysical studies and high-throughput screening. However, conventional imaging and analysis methods only address two-dimensional properties in a wound healing assay, such as gap closure rate. This is unfortunate because biological cells are complex 3D structures, and their dynamics provide significant information about cell physiology. Here, we presented three-dimensional (3D) label-free imaging for wound healing assays and investigated the 3D dynamics of CCM. High-resolution subcellular structures as well as their collective dynamics were imaged and analyzed quantitatively. Our label-free quantitative 3D analysis method provides a unique opportunity to study the behavior of migrating cells during the wound healing process.

## INTRODUCTION

Most biological systems are composed of multiple cells. These collective cell systems involve complex interactions between cells, resulting in a completely different picture compared to single-cell activity (1). Collective cell migration (CCM) includes multiple important phenomena, such as cancer progression, wound healing, and morphogenesis (2). CCM has been widely studied using the wound healing model owing to its simplicity of sample generation and ease of analysis (3). Furthermore, it provides an effective tool for high-throughput screening by observing the effect of various chemical treatments on wound healing speed (4). Numerous studies have observed novel mechanical and molecular interactions between cells during the healing process using this assay (5, 6).

The most common and conventional format of the wound healing model is the two-dimensional (2D) cell monolayer, and the migration of cells is usually imaged using 2D bright-field, phase-contrast, or fluorescence microscopy (7–9). These imaging techniques are utilized to measure the wound healing rate based on size of the wound or the total number of cells inside the initial wound area. Multiple algorithms and software programs have been developed for the analysis of 2D wound healing assay images (10, 11). However, these assay analyses do not consider the three-dimensional (3D) characteristic structures of the cells. The 3D structure and dynamics of subcellular organelles have not been addressed in the context of CCM in a wound healing assay. This is mainly because conventional imaging methods for the study of CCM do not consider 3D subcellular imaging of individual cells.

Here, we presented 3D label-free imaging and quantitative analysis of CCM in a wound healing assay. Exploiting optical diffraction tomography (ODT), a 3D quantitative phase imaging (QPI) technique (12), we demonstrated that refractive index (RI) tomogram measurements reveal the 3D high-resolution structures of individual cells in CCM. By stitching multiple 3D RI tomograms measured at various positions and times, we illustrated long-term and large-scale CCM. We studied both the overall shape and subcellular structures of individual cells using 3D RI information during a time-lapse. Multiple 3D quantities, such as the thickness of cells or RI distribution inside the nuclei in groups of cells with different chemical treatments or wound boundary locations, were explored. This new biophysical approach would readily enable various investigations of the wound healing mechanism, and how the cells react to various chemical signals.

## MATERIALS AND METHODS

### Wound healing assay

NIH3T3 cells (ATCC CRL-1658) were maintained in Dulbecco’s modified Eagle’s medium (DMEM; Gibco) supplemented with 10% fetal bovine serum (FBS; Invitrogen) at 37 °C in a 5% CO_2_ incubator. Approximately 450,000 cells were seeded in TomoDish (Tomocube Inc., Daejeon, Korea), a specially designed cell culture dish to maximize the quality of the tomogram during ODT data acquisition. The TomoDish was coated using 0.01% poly-D-lysine for 15 min and washed 3 times with distilled water. After washing, it was fully dried and stored until use in the experiment. The cells were left to adhere to the dish surface and grow into a confluent monolayer for 28-30 h. The wound was formed by scratch assay using a pipette tip. After scratching, the healing process of the wound was monitored using an ODT microscope. NIH3T3 cells were treated with 1 μg/mL cytochalasin D (Cyto D), which was added to the cell medium at the start of the migration period.

### Optical diffraction tomography

To measure the 3D RI tomograms of cells, we utilized ODT, also known as holotomography (HT). As an optical analogy to X-ray computed tomography, ODT exploits the RI as an intrinsic imaging contrast and reconstructs the 3D RI distribution of unlabeled cells (13–15). Furthermore, ODT does not require the use of exogenous labeling agents or dyes. It not only greatly simplifies the experimental steps but also eliminates obstacles, such as photobleaching or phototoxicity, that originate from the limitations of labeling. This gives ODT an incomparable advantage in monitoring the healing process in 3D for an extended period compared to other imaging techniques.

The ODT measurements of live cells were performed since the cells were maintained at 37 °C and 10% CO_2_ using a live-cell imaging chamber (Tomochamber; Tomocube Inc, Daejeon, South Korea). ODT of wound healing assay cells was performed using a commercial ODT microscope (HT-2H; Tomocube Inc., Daejeon, South Korea). The ODT system used is based on a Mach-Zehnder interferometric microscope equipped with a digital micromirror device (DMD) (16). A coherent laser beam from a diode-pumped solid-state laser (*λ*=532 nm) was used as the illumination source. The beam from the laser is split by a 2×2 single-mode fiber coupler. One beam illuminates a sample as a plane wave, and its incident angle is controlled by projecting time-multiplexed holograms onto the DMD (17). Following this, the diffracted beam for the sample is imaged onto a camera plane, where it generates spatial interference patterns or holograms with a reference laser beam split from the fiber coupler.

For each hologram, both the amplitude and phase images are retrieved using a phase retrieval algorithm (18). Subsequently, from multiple 2D optical holograms of a sample acquired with various illumination angles, the 3D RI distribution of the sample is reconstructed by inversely solving the Helmholtz equation with the Rytov approximation of weak scattering (14). The theoretically calculated lateral and axial spatial resolutions of the optical imaging system used were 110 nm and 360 nm, respectively (19, 20).

Owing to the limited numerical aperture of the condenser and objective lenses, side scattering signals are not collected, which results in the degradation of image quality in reconstructed 3D RI tomograms (21). To address this missing cone problem, an iterative regularization algorithm based on the non-negative constraint was used. Details on the reconstruction procedure and algorithm can be found elsewhere (15, 22).

### Large field-of-view (FoV) imaging

The FoV of each 3D RI tomogram was 82 μm × 82 μm. To perform large FoV measurements, multiple 3D RI tomograms were measured at various lateral positions and stitched into one large FoV tomogram. To accomplish this, a motorized sample stage was synchronized with the image acquisition, and the stitching algorithm was used to seamlessly connect multiple 3D RI tomograms (23).

Multiple field area stitching for wide-field imaging was performed using a stitching function in commercial software (Tomostudio; Tomocube Inc.). The overlap area between each tile was 30% of the area of a single FoV A large area of a sample (up to 1 mm × 1 mm) was imaged with a high resolution (down to 110 nm and 360 nm for the lateral and axial spatial resolutions, respectively) at a high frame rate (up to 3–5 tomograms per s for a FoV of 82 μm × 82 μm). Tomograms were taken every few minutes to hours.

### Cell parameter analysis

3D tomogram imaging by ODT can provide information about the RI in the sample. The volume and thickness of the cells were calculated by thresholding the RI tomogram with respect to the background RI value using a custom-written MATLAB code. The thickness calculated using this method thus directly relates to the thickness of the cell monolayer.

Cell nuclei RI histogram analysis was performed using the ImageJ Analyze-Histogram tool (National Institutes of Health [NIH], Bethesda, MD, USA) after selecting the nuclei region using the freehand selection tool (24). The regions were selected at different depths in the tomogram to obtain volumetric data. The analysis was conducted on 15 cells at the wound boundary (leading cells) and 17 cells that were located at least 8–10 rows of cells away from the boundary (lagging cells).

### Particle image velocimetry (PIV) analysis

PIV analysis was performed using the PIV lab software (25) written in MATLAB. It calculates the velocity information in multiple grid positions from 2D cross-section image pairs. It is based on a discrete Fourier transform technique that calculates the correlation matrix using the Fourier transform of each grid tile and finds the relative position between tiles at different time points. The resulting velocity vectors indicate the change in the cell position as a function of both time and position.

### Statistical analysis

Distributions obtained for RI inside the nuclei of cells in different categories were compared using the two-sample t-test. The shaded error bars in the thickness graph and nuclei RI histogram represent the standard deviation (SD). In the RI distribution graphs, the upper and lower box ranges indicate the 25^th^ and 75^th^ percentiles, respectively.

## RESULTS AND DISCUSSION

### Optical diffraction tomography imaging of the collectively migrating cells

To demonstrate 3D label-free, large-scale FoV imaging of a wound healing assay, the CCM of NIH3T3 fibroblasts was measured (Fig. 1; see Methods). From the 2D multiple holograms measured at various illumination angles, the amplitude and phase images were retrieved using the phase retrieval algorithm (Fig. 1A). Following this, the 3D RI tomogram of a sample was reconstructed using the ODT algorithm. To cover a large FoV, multiple RI tomograms were measured at various lateral positions, from which the 3D RI tomogram with a large FoV was stitched.

**Figure 1.**
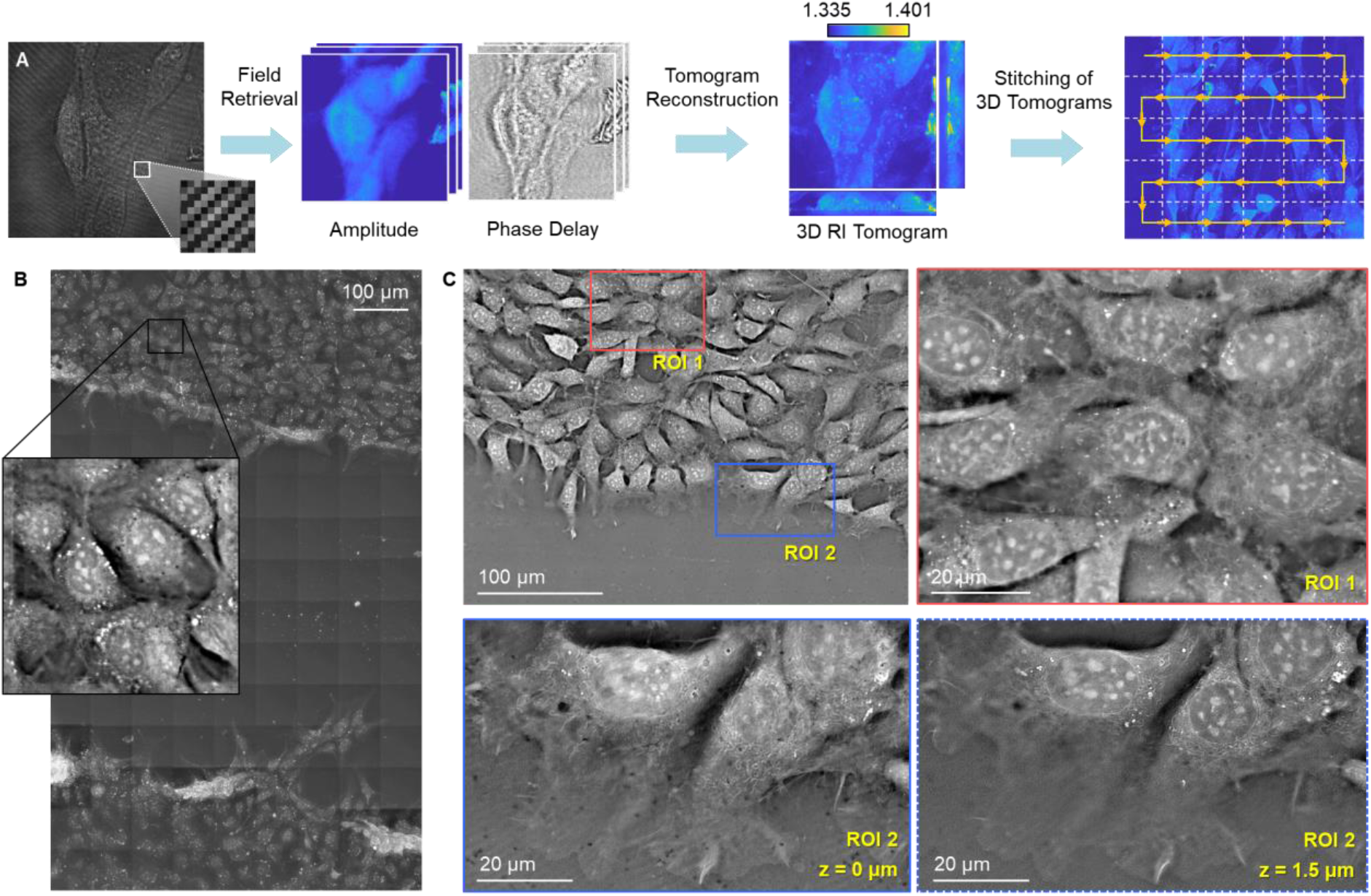
Wide-field imaging of a wound healing assay using optical diffraction tomography (ODT). (A) The reconstruction process of ODT. Multiple holograms are acquired and processed to retrieve the amplitude and phase delay information of the light passing through the sample. The tomogram reconstruction algorithm generates a three-dimensional refractive index (RI) map, which provides quantitative information from the sample. Then, the stitching algorithm based on the correlation method generates a single wide-field RI tomogram using multiple single-field RI tomograms. (B) Maximum intensity projection (MIP) image of a wide-field RI tomogram of a wound healing assay. NIH3T3 cells are collectively migrating into the wound area in the middle, which is an empty space generated by scratching with a pipette tip. (C) A lateral cross-section image of a RI tomogram shows NIH3T3 cells migrating towards the wounded region on the right. The lateral resolution of the imaging system is down to 110 nm, which allows for imaging of various subcellular organelles as shown in the region of interest (ROI) 1. The wound area has no cells, and the cells on the wound boundary form migrating structures as shown in ROI 2. The cell membrane, cytoskeletons, and subcellular organelles all have different morphologies and distributions for different axial positions. Image planes of two different heights are shown for ROI 2.

The reconstructed large-scale RI tomogram of NIH3T3 cells presented large-scale, high-resolution volumetric imaging. Figure 1B is a representative image with the total FoV covering both sides of the wound (670 μm × 1,120 μm). Figure 1C shows the left side of a wound where the cells collectively migrate towards the right. The maximum intensity projection (MIP) along the axial direction visualizes the overall distribution of cells (Fig. 1B).

Unlike conventional imaging methods, HT enables the multiscale tomographic imaging of multicell systems with a large FoV covering both sides of the wound and down to high-resolution subcellular features (Fig. 1C). The cell membrane, nucleus membrane, nucleoli, mitochondria, and vesicles with high RI values were distinctly visualized (regions of interest [ROIs] 1 and 2 in Fig. 1C) without introducing any exogenous labeling. The magnified image of ROI 1 showed the cells that were away from the wound boundary, or so-called lagging cells. Here, the cells were surrounded by other cells, which resulted in a completely different morphology from the leading cells in ROI 2. At z = 0 μm (the bottom of the cell), the complex distribution of cytoskeletons around and at the bottom of the nuclei were clearly visible. The cytoskeletons were also located around the nuclei at z = 1.5 μm, but at this height, they expanded only to the closer peripheral area of the nuclei. Formation of lamellipodia was observed in both cross-sections, showing wrinkled cellular membranes where the cells extended out to the empty wound area. A high RI (bright pixels) was usually observed around and inside the nuclei. These results demonstrate the validity and applicability of HT in the study of CCM.

### Monitoring the thickness of the cell layer

Optical diffraction tomography imaging allows for direct and exact examination of the z-directional thickness of the sample based on 3D quantitative information from the RI. 2D QPI techniques, such as digital holography, have been used for wound healing assays. However, 2D QPI techniques only measure the optical thickness, a coupled parameter between the physical thickness and RI (26, 27), whereas ODT decouples physical thickness from a direct tomographic measurement and provides precise and quantitative thickness information about cells.

The thickness of the migrating cell monolayer was retrieved from a single tomogram as shown in Fig. 2A. The thickness of the region surrounded by a dotted box at multiple time points is shown in Fig. 2B. The lateral position describes the distance from the left end, and each thickness value is calculated from the YZ MIP image. The wound boundary created by scratching the confluent cell layer has regions with some cells that are jammed together. As time passes, these closely packed cells return to the ground. This overall process is shown in the time-lapse thickness data. There initially exists a peak on the frontmost borderline, and then the height gradually decreases with time. In addition, the tendency of leading cells to reach out to the empty wound area and start migration is illustrated by the consistently decreasing slope of the thickness graph line of the frontmost cell.

**Figure 2.**
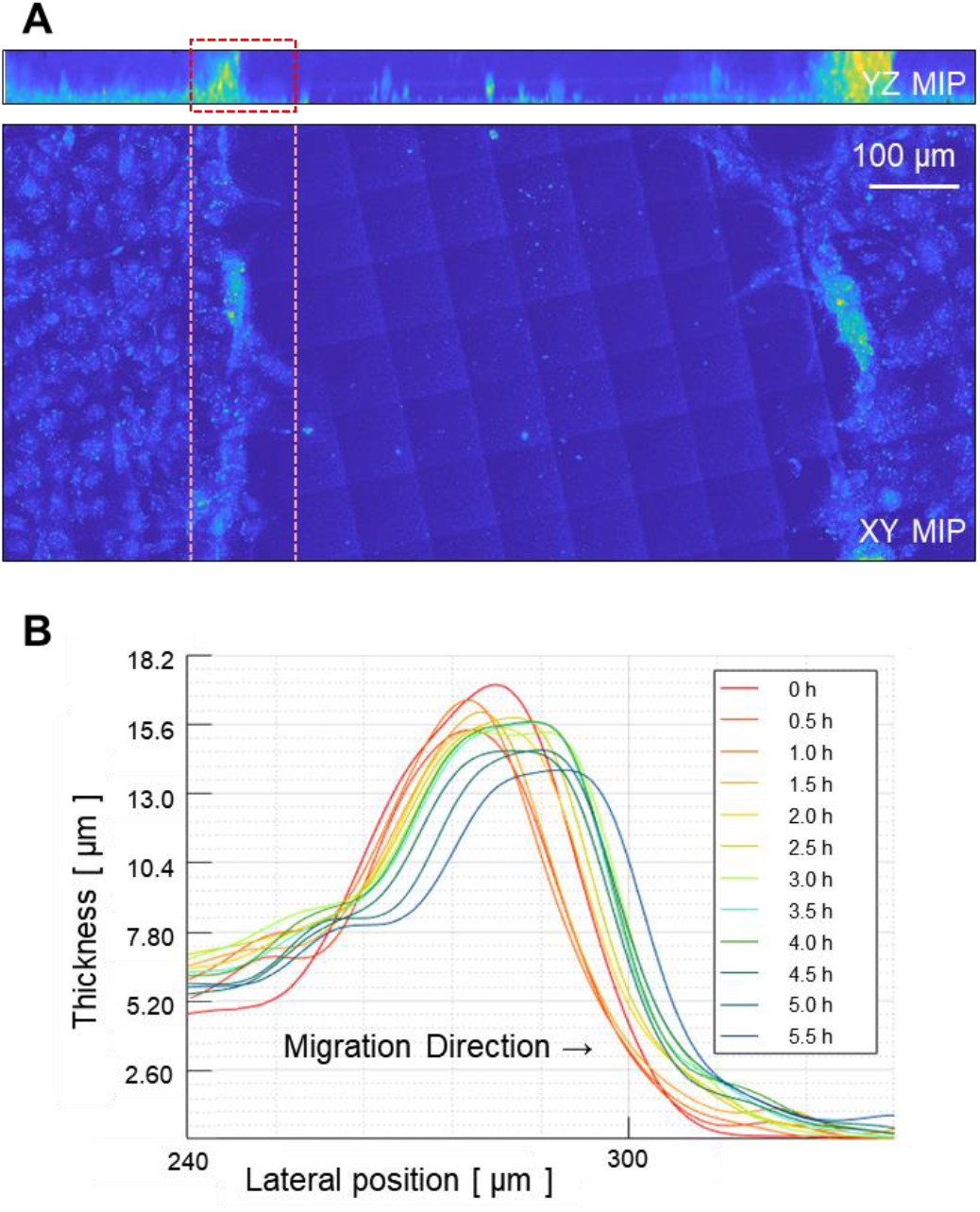
Thickness of the cell monolayer changes over time. (A) Maximum intensity projection (MIP) images from a three-dimensional refractive index tomogram of a wound healing assay. The wound region without any cells is created in the center. The dotted box specifies the boundary region of the wound. Cells inside the dotted box are leading cells, which actively migrate toward the wound area. (B) Thickness value of the cell monolayer is calculated from the YZ MIP image at various time points. Thickness peak point corresponds to the region where multiple cells are jammed together when the wound was created by scratching. Peak value decreases and migrates forward as time passes. Slope in the frontmost region decreases as the cells form migrating structures in the forward direction.

### Quantification of wound healing progress using chemical treatments

To perform 3D tomographic imaging and quantification of migrating cells upon chemical treatment, the physical thickness and 2D migration of individual cells were measured and analyzed simultaneously.

Quantification of wound healing progress or the amount of cell migration can be accomplished using the thickness distribution information of the cell layer in addition to the widely used 2D migration distance information. Fig. 3A shows multiple MIP images of tomograms taken over a time-lapse as the cells migrate towards the empty wound area, which in this case is the upper direction. The control and Cyto D-treated groups were wounded, and their recovery processes were observed. The ODTs are visualized using 2D MIP images, which are similar to the imaging results from other widely used imaging methods, such as bright-field or fluorescence imaging. In addition, ODT can provide 2D images from any point of view using 3D tomographic data. For example, the YZ MIP images on the rightmost side in Fig. 3A demonstrate how the cells are migrating to heal the wound from a unique viewpoint. Based on these data, Fig. 3B shows the migration distance of cells from each group. As expected, the Cyto D-treated group showed a much slower migration speed than the control group.

**Figure 3.**
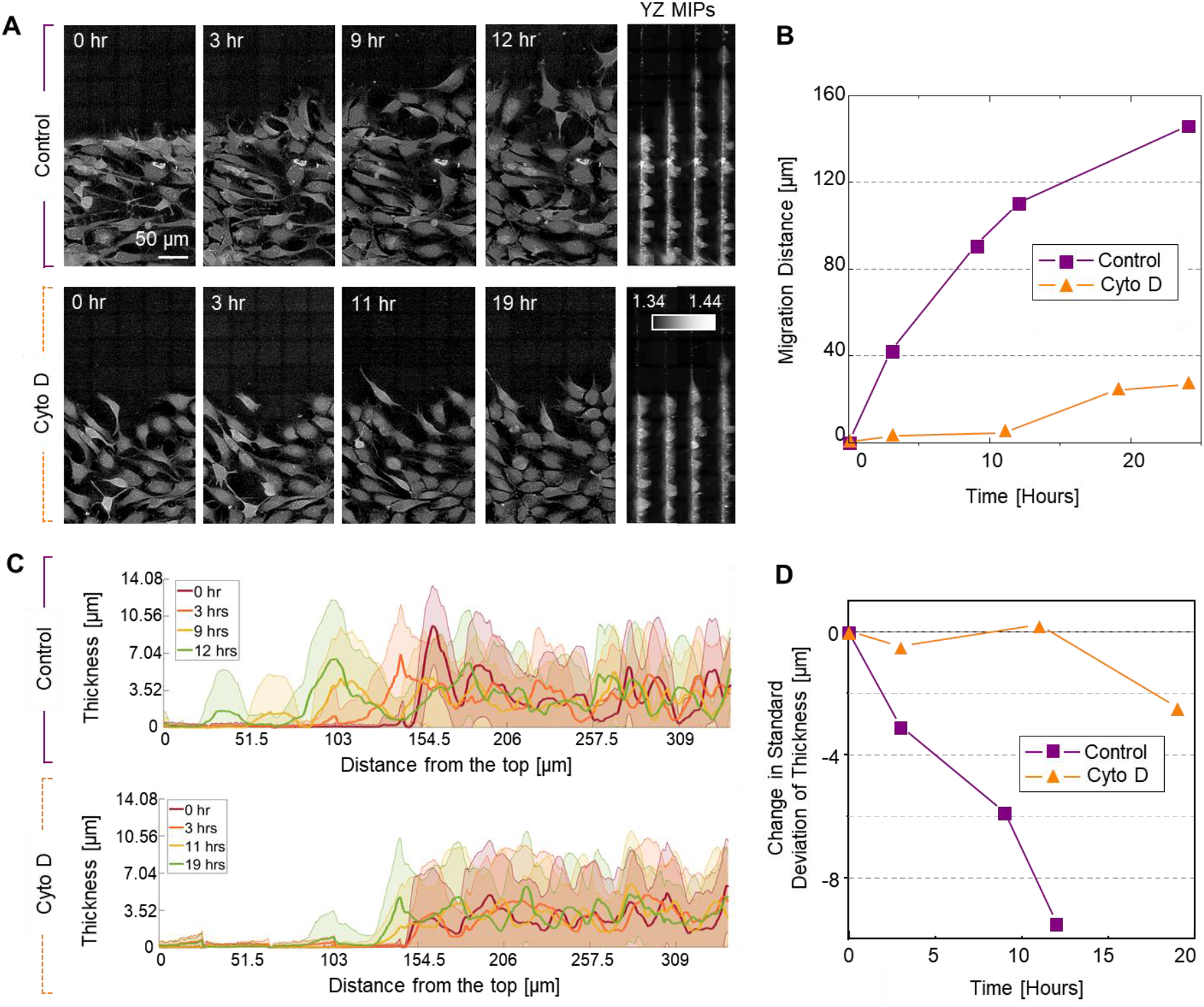
Wound healing process analysis in terms of three-dimensional morphology. (A) Wound healing progress shown in a refractive index tomogram for each control and Cytochalasin D (Cyto D)-treated group of cells. (B) Migration distance is compared for the control and Cyto D-treated cells. (C) Thickness values are calculated and compared. The shaded area represents the standard deviation of the thickness values of the points in the same lateral position. Thickness values in each lateral position do not significantly change after Cyto D treatment. (D) Change in the total standard deviation of thickness values is much smaller for Cyto D-treated cells. Cyto D not only slows the migration speed down, but restricts the thickness variation of the cell layer.

The thickness values were also calculated, and their distributions were studied over time. The average thickness values of the points at the same lateral positions are represented by the solid line and the standard deviation as the shaded area. Thus, the effect of some empty areas between the cells can be reduced and allows for examination of the overall trend of thickness change over time. The standard deviation of all thickness values of each pixel point reflects how much the cells migrate and fill the empty spaces between the cells as well as how much the cells change their shape and become attached to the ground. Over time, the value tends to decrease as shown in Fig. 3D, and the slope is much steeper in the control group compared to the Cyto D-treated group.

### Nuclei of leading and lagging cells exhibit different RI distributions

Several previous studies have been conducted regarding changes in nuclear structure during cell migration. A number of them have revealed that directed cell migration is closely related to the condensation of chromatin fibers and shape of the nucleus (28–30). In addition, it has been determined that the ratio of euchromatin to heterochromatin is a factor in determining nuclear stiffness and affects cell migration, which requires further investigation (31–33). Other studies identified a link between the nucleoli and the extracellular force applied to the cell (34) as well as the role of A-type lamin for sustaining directed cell migration (35)

To demonstrate the capability of the present method, the nuclei of leading cells actively migrating towards the wound boundary and lagging cells surrounded by other cells far behind the wound boundary were compared. In Fig. 4A, representative images of lagging and leading cell nuclei are shown as yellow solid circles and red dotted circles, respectively. The RI of a cell and its nuclei are strongly inhomogeneous (Fig. 4B). Higher RIs exist mainly inside or near the nucleus of a cell. For further investigation, multimodal ODT and fluorescence imaging were conducted to confirm the range of the RI distributions, which correlated well with some of the well-known molecular contents inside the nucleus. Hoechst dye, a blue fluorescent dye widely used for labeling DNA (36), showed a good correlation with the RI values from 1.364 to 1.373.

**Figure 4.**
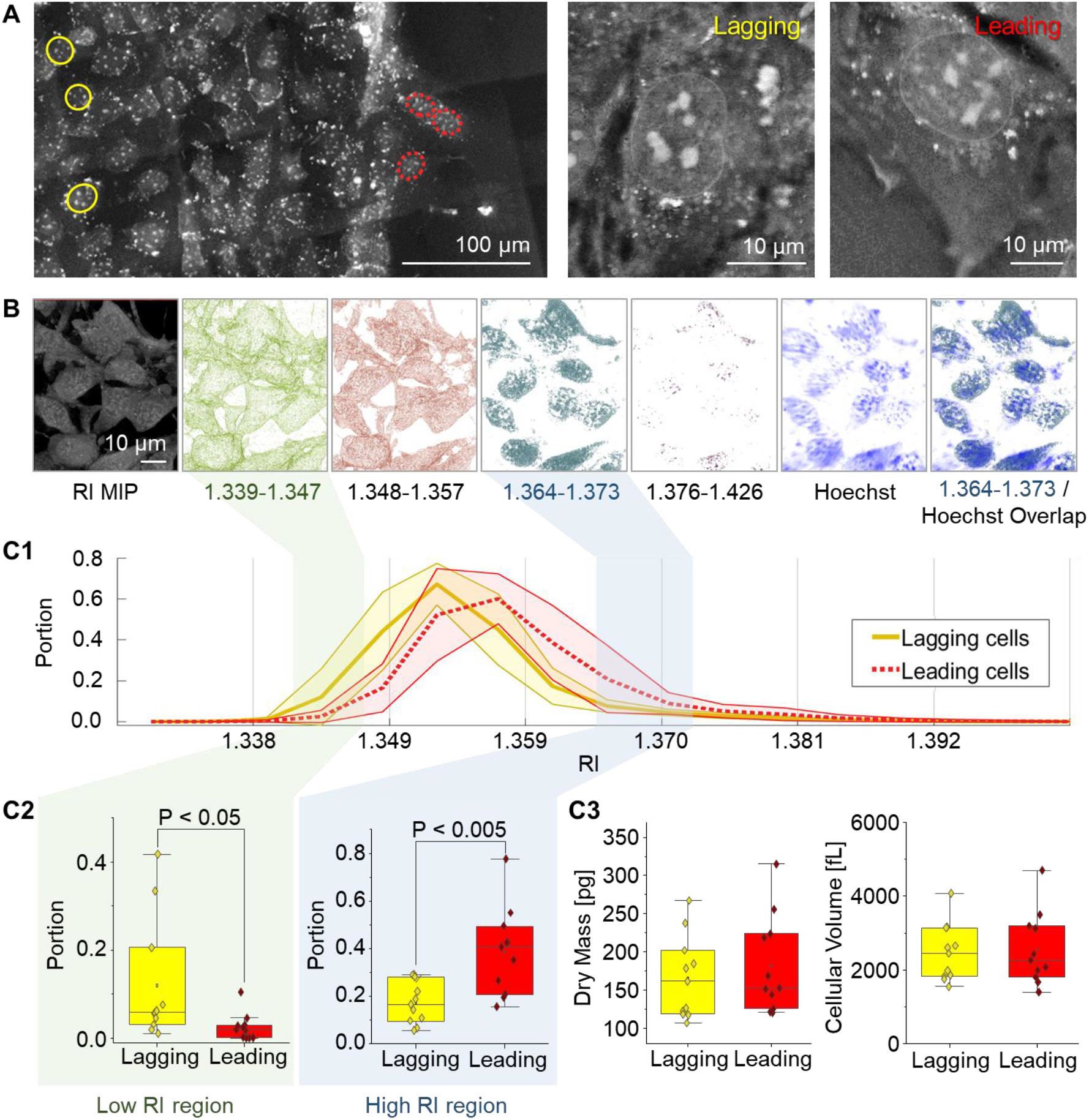
Refractive index (RI) distribution inside the nuclei of cells near and away from the wound boundary. (A) Yellow solid circles represent the nuclei of lagging cells, whereas red dotted circles represent the nuclei of leading cells. The nuclear membrane is visible as a bright, thin circular shape in each cell. (B) RI tomogram visualization using different-colored voxels within certain RI ranges. The regions with RI values from 1.364 to 1.373 and the Hoechst fluorescence signal distributions overlap. (C1) RI histogram inside the volume of each nucleus within lagging and leading cells. (C2) Lagging cells have a higher portion of RI values in the low RI region than leading cells. On the contrary, leading cells have a higher portion of RI values in the high RI region than lagging cells. (C3) Cellular dry mass and cellular volume were calculated and compared between the lagging and leading cells. There is no meaningful difference between the two groups.

For systematic comparison, the RI histograms inside the 3D volume of nuclei in lagging and leading cells were compared (Fig. 4C1). The lagging and leading cells had opposite dominance in the low and high RI regions inside their nuclei (Fig. 4C2). The results showed that a higher RI range was included in the RI range that showed a good correlation with the Hoechst signal (1.364-1.373). The data demonstrated that the leading cells had more regions in their nuclei with higher RI values, which correlated well with the fluorescence signal labeling DNA. The RI values provide quantitative information on how optically densely the molecules are packed together. The dry mass, which is the total amount of molecules that exist inside each cell, and the volume of each cell revealed a negligible difference in the groups as shown in Fig. 4C3. This data implies that leading cells have more densely packed DNA in their nuclei than lagging cells.

Overall, the aforementioned results suggest that the leading cells in directed migration motion have more condensed contents, including DNA, which constitute the higher RI regions inside their nuclei. This is further evidence that environmental cues alter the morphology of the nuclei and are related to the way cells react to them. Although RI tomograms do not generally have molecular specificity when compared to fluorescence or other labeling methods, they provide quantitative information about the whole nuclei without missing other unlabeled molecules. In addition, the RI values directly relate to how densely molecules are packed compared to previous studies that were necessary to find a quantity that indirectly represents the level of condensation of certain molecules. Some studies have proposed indices that are calculated indirectly based on thresholding or edge detection in the fluorescence image, but they fundamentally depend on the intensity of the fluorescence signals in the image (37, 38).

### Multilayer particle image velocimetry

To further demonstrate the potential of the present approach, we analyzed the collective motions of cells at various axial positions using PIV (Fig. 5). The boundary where the cells protrude from their original position to the empty area was examined at various depths. Fig. 5A shows the results in which the green vectors indicate the migration velocity at each grid point and the yellow lines are the streamlines. From 0.5-1 h after scratching, most of the cells remained, except for the leading cells. The leading cells clearly showed the largest migration velocity, and their movement was larger near the bottom coverslip surface. The migration velocity and its directionality toward the wound area become larger as time passed, making the streamline perpendicular to the wound edge. Fig. 5B represents the average map of all velocity vector map results at different time steps on the plane with a height of z = 2.34 μm. The point where all of the average streamlines start was where the highest number of cells per volume was located. This illustrated the pushing forces in the compact region between adjacent cells in addition to the different dynamics of leading and lagging cells. The migration velocity vectors (Fig. 5C) demonstrated an asymmetric distribution concerning the origin pointing toward the migration direction.

**Figure 5.**
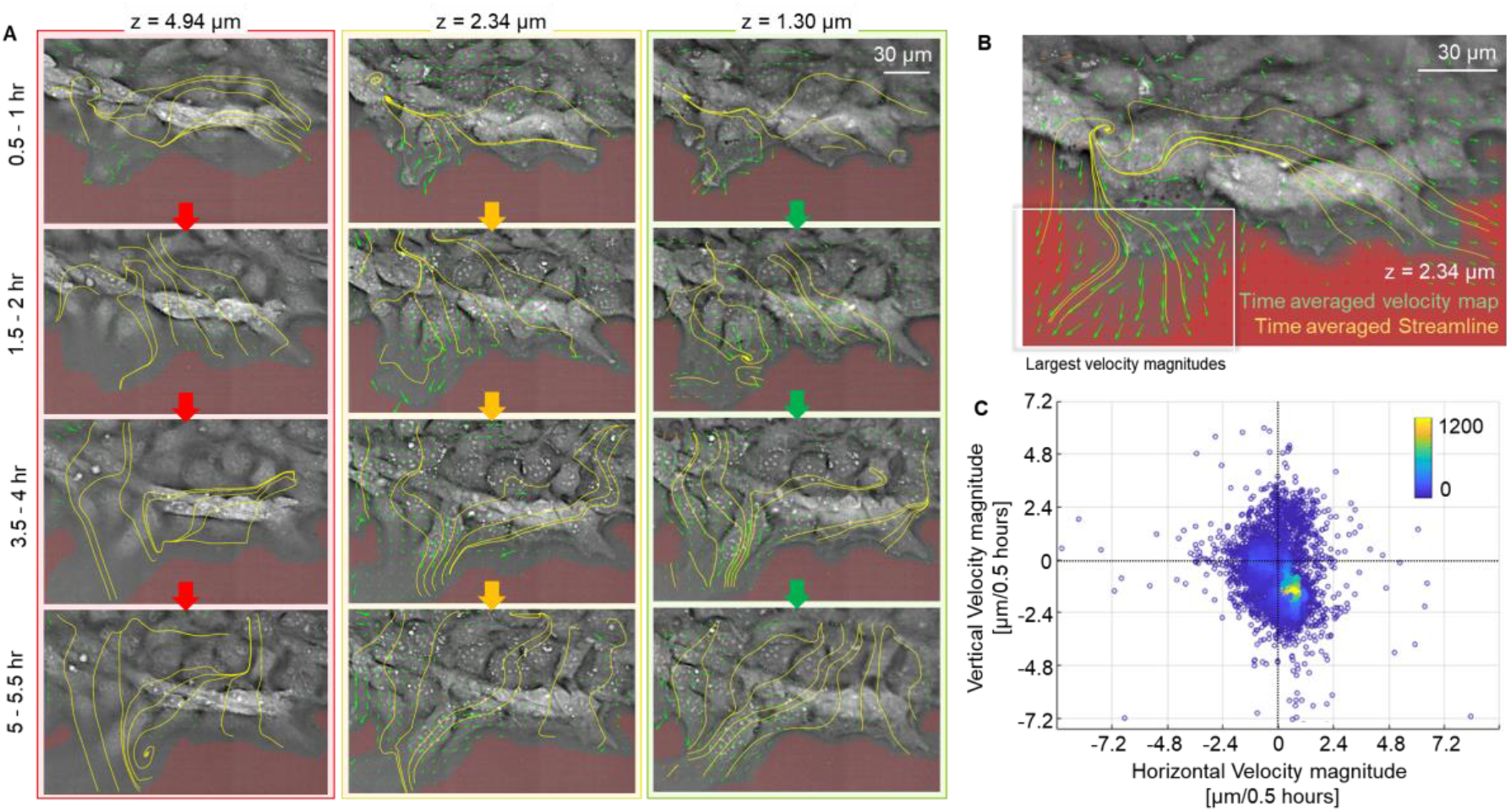
Particle image velocimetry (PIV) analysis on cell migration using optical diffraction tomography. (A) PIV analysis of different planes of tomograms at various timepoints. Each green arrow shows the migration velocity at each grid point, and the yellow lines are the streamlines. They differ both temporally and spatially. Velocity magnitudes are generally larger in the lower part of the cells that are nearer to the wound boundary. The streamlines are initially aligned orthogonal to the migration direction but gradually become parallel over time in all planes. (B) Average velocity map and average streamline for all timepoints at z = 2.34 μm. The leader cell is characterized as the cell with the largest velocity magnitudes on the map. (C) The velocity vectors are plotted in a single graph. The vertical velocity distribution is asymmetric with respect to zero, showing that the cells are migrating toward the wound area in total.

## CONCLUSION

To perform 3D label-free imaging and quantification of a wound healing assay, ODT imaging, synchronization of a motorized sample stage, and a stitching algorithm were combined. With the present approach, we investigated the wound healing mechanism in confluent monolayer NIH3T3 cells. This method enables the measurement of various biophysical parameters in a wound healing assay, which are inaccessible or challenging to access with conventional methods. We visualized and quantified individual migrating cells and their subcellular organelles. The physical thicknesses were directly retrieved from the tomographic measurements. Migration of cells at multiple depth positions was proven, which clearly demonstrates that cellular migration and associated subcellular movements in a wound healing model exhibit 3D characteristics, even for monolayered cells. Furthermore, the time-lapse measurement of the thickness of a confluent monolayer from the 3D tomogram data showed the effects of Cyto D treatment. Finally, the RIs inside the nuclei were distributed differently in leading cells compared to lagging cells.

Thus far, 2D QPI techniques, such as digital holographic microscopy, have been applied to the study of cell migration or wound healing assays (39–42). Although the use of 2D QPI techniques has included label-free and quantitative imaging capabilities, only limited 2D optical thickness information can be measured, and the 3D morphology of individual cells and subcellular structures is not accessible.

In this study, we presented label-free tomographic imaging and quantification of a wound healing assay and illustrated the migration of NIH3T3 cells. We also demonstrated the proof of principle and expect that the current approach can be further exploited in various biophysical studies to stimulate new research approaches. For example, tomographic subcellular information in a migration assay will also promote the development of the theoretical modeling of wound healing (43). In addition, as scientists’ interest in the cell nucleus rotation or its morphology during cell migration increases, further examination of the RI inside the nuclei and other analyses become possible using the ODT wound healing data accompanied by quantitative analysis using the dry mass calculation from the raw data. Furthermore, our work was limited to 2D wound healing assays, but 2.5D or 3D wound healing models are readily accessible for examination using ODT. How cells migrate in a 3D extracellular matrix can be directly imaged without labeling for a very long period. Finally, 3D PIV analysis combined with a 3D phase correlation algorithm would allow direct visualization of how leading cells protrude their pseudopods and move towards the empty area.

Our method would also be of particular interest to the pharmaceutical industry for drug discovery. It has a unique ability to examine 3D dynamics over a wide FoV for a long period of time and without any labeling. This provides an opportunity to study and analyze wound healing mechanisms using a new viewpoint. Studying wound healing models in a 3D environment by applying this technology will allow for a better and more systematic understanding of CCM as well as the wound healing mechanism.

## AUTHOR CONTRIBUTIONS

A.J.L. and Y.P. conceived the initial idea. A. J. L. performed the experiments and analyzed the data. H.H. and W.S.P. provided the experimental and analysis methods. All authors wrote and revised the manuscript. Correspondence and requests for materials should be addressed to Y.P.

## ACKNOWLEDGMENTS

This work was supported by KAIST, BK21+ program, Tomocube, and National Research Foundation of Korea (2017M3C1A3013923, 2015R1A3A2066550, 2018K000396).

## REFERENCES

1. Sadati, M., N. Taheri Qazvini, R. Krishnan, C. Y. Park, and J. J. Fredberg. 2013. Collective migration and cell jamming. Differentiation 86(3):121–125.

2. Friedl, P., and D. Gilmour. 2009. Collective cell migration in morphogenesis, regeneration and cancer. Nature Reviews Molecular Cell Biology 10(7):445–457.

3. Grada, A., M. Otero-Vinas, F. Prieto-Castrillo, Z. Obagi, and V. Falanga. 2017. Research Techniques Made Simple: Analysis of Collective Cell Migration Using the Wound Healing Assay. Journal of Investigative Dermatology 137(2):e11–e16.

4. Muniandy, K., S. Gothai, W. S. Tan, S. S. Kumar, N. Mohd Esa, G. Chandramohan, K. S. Al-Numair, and P. Arulselvan. 2018. In Vitro Wound Healing Potential of Stem Extract of Alternanthera sessilis. Evid Based Complement Alternat Med 2018:3142073–3142073.

5. Riahi, R., J. Sun, S. Wang, M. Long, D. D. Zhang, and P. K. Wong. 2015. Notch1–Dll4 signalling and mechanical force regulate leader cell formation during collective cell migration. Nature Communications 6(1):6556.

6. Plutoni, C., E. Bazellieres, M. Le Borgne-Rochet, F. Comunale, A. Brugues, M. Séveno, D. Planchon, S. Thuault, N. Morin, S. Bodin, X. Trepat, and C. Gauthier-Rouvière. 2016. P-cadherin promotes collective cell migration via a Cdc42-mediated increase in mechanical forces. The Journal of Cell Biology 212(2):199–217.

7. Stamm, A., K. Reimers, S. Strauß, P. Vogt, T. Scheper, and I. Pepelanova. 2016. In vitro wound healing assays - State of the art. BioNanoMaterials 0.

8. Liang, C.-C., A. Y. Park, and J.-L. Guan. 2007. In vitro scratch assay: a convenient and inexpensive method for analysis of cell migration in vitro. Nature Protocols 2(2):329–333.

9. Yue, P. Y. K., E. P. Y. Leung, N. K. Mak, and R. N. S. Wong. 2010. A Simplified Method for Quantifying Cell Migration/Wound Healing in 96-Well Plates. Journal of Biomolecular Screening 15(4):427–433.

10. Zaritsky, A., S. Natan, J. Horev, I. Hecht, L. Wolf, E. Ben-Jacob, and I. Tsarfaty. 2011. Cell motility dynamics: a novel segmentation algorithm to quantify multi-cellular bright field microscopy images. PLoS One 6(11):e27593–e27593.

11. Gebäck, T., M. M. P. Schulz, P. Koumoutsakos, and M. Detmar. 2009. TScratch: a novel and simple software tool for automated analysis of monolayer wound healing assays. BioTechniques 46(4):265–274.

12. Park, Y., C. Depeursinge, and G. Popescu. 2018. Quantitative phase imaging in biomedicine. Nat Photonics 12(10):578–589.

13. Kim, D., S. Lee, M. Lee, J. Oh, S.-A. Yang, and Y. Park. 2017. Refractive index as an intrinsic imaging contrast for 3-D label-free live cell imaging. BioRxiv:106328.

14. Wolf, E. 1969. Three-dimensional structure determination of semi-transparent objects from holographic data. Optics communications 1(4):153–156.

15. Kim, K., J. Yoon, S. Shin, S. Lee, and S.-A. Yang. 2016. Optical diffraction tomography techniques for the study of cell pathophysiology. Journal of Biomedical Photonics Engineering 2(2).

16. Shin, S., K. Kim, T. Kim, J. Yoon, K. Hong, J. Park, and Y. Park. 2016. Optical diffraction tomography using a digital micromirror device for stable measurements of 4D refractive index tomography of cells. In Quantitative Phase Imaging II. International Society for Optics and Photonics. 971814.

17. Shin, S., K. Kim, J. Yoon, and Y. Park. 2015. Active illumination using a digital micromirror device for quantitative phase imaging. Opt Lett 40(22):5407–5410.

18. Debnath, S. K., and Y. Park. 2011. Real-time quantitative phase imaging with a spatial phase-shifting algorithm. Optics letters 36(23):4677–4679.

19. Lauer, V. 2002. New approach to optical diffraction tomography yielding a vector equation of diffraction tomography and a novel tomographic microscope. Journal of Microscopy 205(2):165–176.

20. Park, C., S. Shin, and Y. Park. 2018. Generalized quantification of three-dimensional resolution in optical diffraction tomography using the projection of maximal spatial bandwidths. JOSA A 35(11):1891–1898.

21. Lim, J., K. Lee, K. H. Jin, S. Shin, S. Lee, Y. Park, and J. C. Ye. 2015. Comparative study of iterative reconstruction algorithms for missing cone problems in optical diffraction tomography. Optics express 23(13):16933–16948.

22. Kim, K., H. Yoon, M. Diez-Silva, M. Dao, R. R. Dasari, and Y. Park. 2013. High-resolution three-dimensional imaging of red blood cells parasitized by Plasmodium falciparum and in situ hemozoin crystals using optical diffraction tomography. Journal of biomedical optics 19(1):011005.

23. Hugonnet, H., Y. W. Kim, M. Lee, S. Shin, R. H. Hruband, S. Hong, and Y. Park. 2020. Multiscale label-free volumetric holographic histopathology of thick-tissue slides with subcellular resolution. BioRxiv:2020.2007.2015.205633.

24. Schindelin, J., I. Arganda-Carreras, E. Frise, V. Kaynig, M. Longair, T. Pietzsch, S. Preibisch, C. Rueden, S. Saalfeld, B. Schmid, J.-Y. Tinevez, D. J. White, V. Hartenstein, K. Eliceiri, P. Tomancak, and A. Cardona. 2012. Fiji: an open-source platform for biological-image analysis. Nature Methods 9(7):676–682.

25. Thielicke, W., and E. Stamhuis. 2014. PIVlab – Towards User-friendly, Affordable and Accurate Digital Particle Image Velocimetry in MATLAB. Journal of Open Research Software 2.

26. Bettenworth, D., P. Lenz, P. Krausewitz, M. Brückner, S. Ketelhut, D. Domagk, and B. Kemper. 2014. Quantitative stain-free and continuous multimodal monitoring of wound healing in vitro with digital holographic microscopy. PLoS One 9(9).

27. Bettenworth, D., A. Bokemeyer, C. Poremba, N. S. Ding, S. Ketelhut, P. Lenz, and B. Kemper. 2018. Quantitative phase microscopy for evaluation of intestinal inflammation and wound healing utilizing label-free biophysical markers. Histol. Histopathol 33:417–432.

28. Gerlitz, G., I. Livnat, C. Ziv, O. Yarden, M. Bustin, and O. Reiner. 2007. Migration cues induce chromatin alterations. Traffic 8(11):1521–1529.

29. Gerlitz, G., and M. Bustin. 2011. The role of chromatin structure in cell migration. Trends in Cell Biology 21(1):6–11.

30. Gerlitz, G., and M. Bustin. 2010. Efficient cell migration requires global chromatin condensation. Journal of Cell Science 123(13):2207–2217.

31. Calero-Cuenca, F. J., C. S. Janota, and E. R. Gomes. 2018. Dealing with the nucleus during cell migration. Current Opinion in Cell Biology 50:35–41.

32. Pajerowski, J. D., K. N. Dahl, F. L. Zhong, P. J. Sammak, and D. E. Discher. 2007. Physical plasticity of the nucleus in stem cell differentiation. Proceedings of the National Academy of Sciences 104(40):15619.

33. Erdel, F., M. Baum, and K. Rippe. 2015. The viscoelastic properties of chromatin and the nucleoplasm revealed by scale-dependent protein mobility. Journal of Physics: Condensed Matter 27(6):064115.

34. Maniotis, A. J., C. S. Chen, and D. E. Ingber. 1997. Demonstration of mechanical connections between integrins, cytoskeletal filaments, and nucleoplasm that stabilize nuclear structure. Proceedings of the National Academy of Sciences 94(3):849–854.

35. Houben, F., C. H. M. P. Willems, I. L. J. Declercq, K. Hochstenbach, M. A. Kamps, L. H. E. H. Snoeckx, F. C. S. Ramaekers, and J. L. V. Broers. 2009. Disturbed nuclear orientation and cellular migration in A-type lamin deficient cells. Biochimica et Biophysica Acta (BBA) - Molecular Cell Research 1793(2):312–324.

36. Bucevicius, J., G. Lukinavicius, and R. Gerasimaite. 2018. The Use of Hoechst Dyes for DNA Staining and beyond. Chemosensors 6:18.

37. Sosnik, J., W. A. Vieira, K. A. Webster, K. R. Siegfried, and C. D. McCusker. 2017. A new and improved algorithm for the quantification of chromatin condensation from microscopic data shows decreased chromatin condensation in regenerating axolotl limb cells. PLoS One 12(10):e0185292.

38. Irianto, J., D. A. Lee, and M. M. Knight. 2014. Quantification of chromatin condensation level by image processing. Medical Engineering & Physics 36(3):412–417.

39. Tolde, O., A. Gandalovičová, A. Křížová, P. Veselý, R. Chmelík, D. Rosel, and J. Brábek. 2018. Quantitative phase imaging unravels new insight into dynamics of mesenchymal and amoeboid cancer cell invasion. Scientific reports 8(1):1–13.

40. Guo, P., J. Huang, and M. A. Moses. 2018. Quantitative phase imaging characterization of tumor-associated blood vessel formation on a chip. In Quantitative Phase Imaging IV. International Society for Optics and Photonics. 105031O.

41. Langehanenberg, P., L. Ivanova, I. Bernhardt, S. Ketelhut, A. Vollmer, D. Dirksen, G. K. Georgiev, G. von Bally, and B. Kemper. 2009. Automated three-dimensional tracking of living cells by digital holographic microscopy. Journal of biomedical optics 14(1):014018.

42. Rezaei, M., J. Cao, K. Friedrich, B. Kemper, O. Brendel, M. Grosser, M. Adrian, G. Baretton, G. Breier, and H.-J. Schnittler. 2018. The expression of VE-cadherin in breast cancer cells modulates cell dynamics as a function of tumor differentiation and promotes tumor–endothelial cell interactions. Histochemistry and cell biology 149(1):15–30.

43. Arciero, J. C., Q. Mi, M. F. Branca, D. J. Hackam, and D. Swigon. 2011. Continuum model of collective cell migration in wound healing and colony expansion. Biophys J 100(3):535–543.

